# *CXCR5* gene expression in human lymph node CD8^+^ T cells is regulated by DNA methylation and nucleosomal occupancy

**DOI:** 10.1101/2020.07.17.208470

**Authors:** Funsho J. Ogunshola, Werner Smidt, Anneta F. Naidoo, Thandeka Nkosi, Thandekile Ngubane, Trevor Khaba, Omolara O. Baiyegunhi, Sam Rasehlo, Ismail Jajbhay, Krista L. Dong, Veron Ramsuran, Johan Pansegrouw, Thumbi Ndung’u, Bruce D. Walker, Tulio de Oliveria, Zaza M. Ndhlovu

## Abstract

CD8^+^ T cells play an important role in viral and tumour control. However, in human lymph nodes (LNs), only a small subset of CD8^+^ T cells called follicular CD8^+^ T cells (fCD8s) expresses CXCR5, the chemokine receptor required for cell migration into B cell follicles, thought to promote immune evasion. Here we obtained LNs from HIV infected persons to investigate regulation of CXCR5 expression in lymphoid CD8^+^ T cells, and compared this to the more abundant CXCR5 expressing T follicular CD4^+^ helper cells (GCTfh). Our results show that DNA hypermethylation and closed chromatin at the transcriptional start site (TSS) prevent CXCR5 expression in non-fCD8s. We also found that greater nucleosomal density at the CXCR5 TSS could be responsible for reduced CXCR5 expression in fCD8s relative to GCTfh. Together, these data provide critical insights into both the underlying molecular mechanisms that repress CXCR5 expression in non-fCD8s and the plausible mechanism responsible for the low CXCR5 expression in fCD8s, with implications for HIV cure strategies.

**Author Summary:** A paucity of CD8^+^ T cells that express CXCR5, the chemokine receptor critical for entering the B cell follicles of secondary lymphoid tissues have recently been described. Animal studies have revealed transcriptional networks that govern the expression of CXCR5 in CD8^+^ T cells. However, it is not known if similar or additional networks regulate the expression of CXCR5 in human CD8^+^ T cells. In this study, we demonstrated that DNA methylation coupled with chromatin compaction at the transcriptional start site (TSS) of *CXCR5* gene prevent the expression CXCR5 in human CD8^+^ T cells. In addition, we observed greater nucleosomal occupancy at the TSS of *CXCR5* gene which could impact expression levels of CXCR5 in human CXCR5^+^CD8^+^ T cells. This study revealed multitiered epigenetic mechanisms that repress CXCR5 expression in human CD8^+^ T cells, with implications for HIV cure strategy or eradication of B cell-derived tumours.

## Introduction

Upon infection, viral antigens prime naïve CD8^+^ T cells in secondary lymphoid tissues to differentiate into effector cytotoxic CD8^+^ T cells and migrate to sites of infection, guided by chemokine-chemokine receptor interactions (1). In the case of human immunodeficiency virus (HIV) infection, secondary lymphoid tissues serve as the major site of replication (2–4); and germinal centers (GCs) in the B cell follicles of LN serve as major sites of HIV persistence during suppressive antiretroviral therapy (ART) (5–7). CXCR5 expression facilitates direct trafficking of T cells to GCs by sensing CXCL13 producing cells, which reside within LNs (8–10). However, CD8^+^ T cells typically lack CXCR5 expression and are therefore generally excluded from B cell follicles within LN (11, 12) which is thought to be partially responsible for HIV persistence in this compartment, particularly during ART (13, 14). Similar mechanisms contribute to persistence of tumours in lymphoid tissues (15). Thus, development of novel strategies for boosting pathogen-specific CD8^+^ T cell migration to B cell follicles could enhance immune clearance of HIV infected cells and tumour cells such as B cell lymphomas.

A small subset CXCR5 expressing CD8^+^ T cells called follicular CD8^+^ T cells (fCD8) has recently been described, to have the capacity to infiltrate B cell follicles and eliminate HIV infected cells or tumour cells (16–18). Human and animal studies have shown that the frequency of fCD8s inversely correlates with HIV or simian immunodeficiency virus (SIV) viral load (16, 19, 20), suggesting that increased infiltration of fCD8s in B cell follicles can result in enhanced immune control. Indeed, some studies demonstrate direct anti-HIV activity of fCD8s (16, 17). In addition, in the SIV model, CD8^+^ T depletion is associated with modest increase of SIV infected cells in B cell follicles (12), suggesting their involvement in mediating control of virus replication in the follicles. Moreover, in follicular lymphoma (FL), the second most frequent B-Cell lymphoma in adults (21), increased infiltration of CD8^+^ T cells into B cell follicles is associated with improved disease prognosis (18). Thus, detailed understanding of the regulatory mechanisms that govern the expression of CXCR5, the chemokine receptor required for CD8^+^ T cells migration to B cell follicles is highly relevant to the development of curative strategies for HIV and B cell lymphomas (18).

Animal studies have attempted to define the transcriptional regulatory networks that distinguish fCD8s from non-fCD8s. These studies have implicated a number of transcriptional factors (TFs) including B lymphocyte-induced maturation protein-1 (Blimp1) and B-cell lymphoma 6 protein (BCL6) coupled with T-cell factor 1 (TCF1), and inhibitor DNA binding 2 and 3 (Id2 and Id3), which together form a transcriptional circuit that govern fCD8 differentiation (16, 22). Additionally, *in vitro* stimulation of CD8^+^ T cells from rhesus macaques with inflammatory cytokines such as TGF-β, IL-12 and IL-23 promotes fCD8 differentiation (23). Together, these studies provide an important framework for potential regulatory networks. However, the underlying molecular processes that govern fCD8 differentiation remain largely unknown. Moreover, it is not yet clear how these animal studies translate to human diseases.

Here we report a detailed investigation of the epigenetic and transcriptional processes that regulate *CXCR5* gene expression in human CD8^+^ T cells. We test the hypothesis that epigenetic mechanisms, acting in conjunction with specific transcription factors, play a critical role in regulating CXCR5 expression on human CD8^+^ T cells (24). This hypothesis is based on the premise that epigenetic mechanisms such as DNA methylation, chromatin state and accessibility influence gene expression during cell differentiation and maturation (24–26). Furthermore, the density and positioning of nucleosomes around the genomic DNA can regulate the levels of a gene expression by modulating DNA accessibility to TFs (27).

To test this hypothesis, we investigated *CXCR5* gene regulation in lymphoid CD8^+^ T cells in the setting of HIV infection using DNA bisulfite sequencing in combination with the Assay for Transposase-Accessible Chromatin using Sequencing (ATAC-Seq) and RNA-Seq. We found that DNA methylation and chromatin conformation regulate *CXCR5* in human CD8^+^ T cells. Computational analysis further revealed nucleosomal occupancy and positioning around the TSS of the *CXCR5* gene as a plausible mechanism involved in limiting the expression of CXCR5 in fCD8s. This study reveals epigenetic processes that play a pivotal role in limiting the expression of CXCR5 in human CD8^+^ T cells. These results could be the basis for rationale development of novel strategies for increasing CD8^+^ T cells trafficking into B cell follicles where they are needed to clear pathogens such as HIV and B cell lymphomas.

## Results

### Description of study samples and design

This study included 17 participants from the FRESH (Female Rising through Education, Support and Health) program, a socioeconomic and HIV prevention intervention for HIV uninfected women at high risk of infection in KwaZulu Natal, South Africa, designed to facilitate identification of hyperacute infection (28). Participants were classified into 3 groups. Group 1 consisted of 5 HIV negative participants. Group 2 included 7 HIV infected individuals who were on ART for >1 year and were fully suppressed at the time of sample collection. Group 3 included 5 individuals with untreated HIV infection for >1 year with median viral load of 15,068 copies/ml at the time of sample collection. Subjects and time-points were chosen based on sample availability. The clinical characteristics of the study participants are summarized in **Table 1**.

**Table 1.**
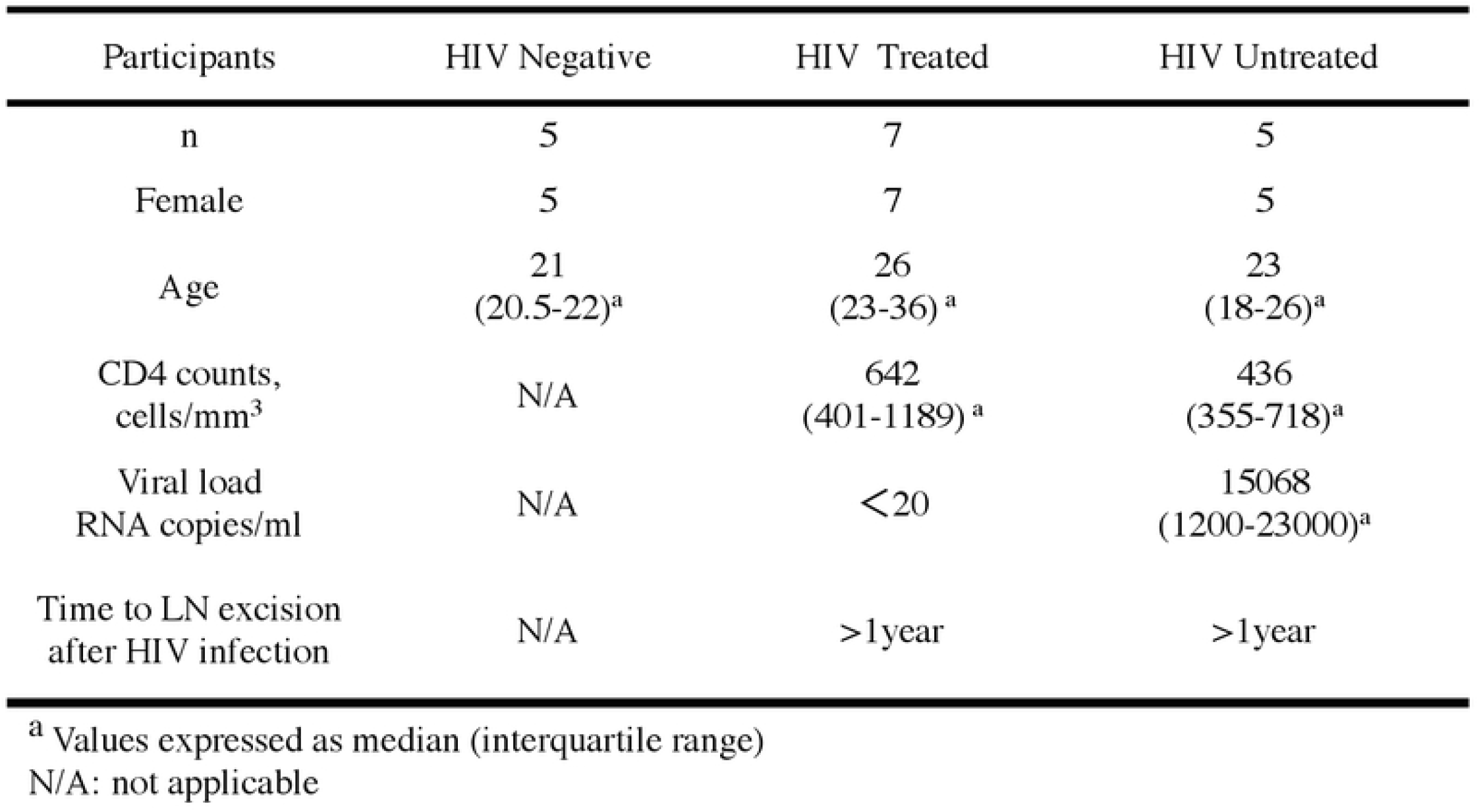
Demographic and clinical characteristics of the study participants

To address our hypothesis, we conducted a series of experiments using one excisional LN and paired peripheral blood sample per study participant. We began by performing flow cytometry on all 17 LN samples to establish the frequency of fCD8s in each experimental group. This was followed by image analysis of fixed LN tissue samples from 9 donors (3 from each experimental group). Imaging studies were used to substantiate the flow data and to determine the localization of CD8^+^ T cell subsets within LNs in health and in HIV disease. A subset of 5 HIV infected participants (3 HIV treated and 2 untreated) were then selected based on sample availability and used for mechanistic studies to define epigenetic processes and transcriptional factors that regulate *CXCR5* gene expression in human CD8^+^ T cells. Details of the experimental design and samples used for each sub study are summarized in the flowchart and cartoon depicted in **supplementary Fig. 1A and B**.

### Phenotypic characterization of fCD8s in HIV infected subjects

Recently, fCD8s were described as tissue resident CD8^+^ T cells (31). To assess whether CD8^+^ T cells that have the follicular-homing phenotype (fCD8s) were indeed localized in the lymphoid tissues during HIV infection, we first used flow cytometry to measure the frequency of fCD8s in LN and in peripheral blood mononuclear cells (PBMCs) in all 3 study groups. Consistent with a recent study (31), we observed a significantly higher frequency of fCD8s in LN compared to PBMCs in all the groups (p<0.0001) **(Fig. 1A)**.

**Figure 1:**
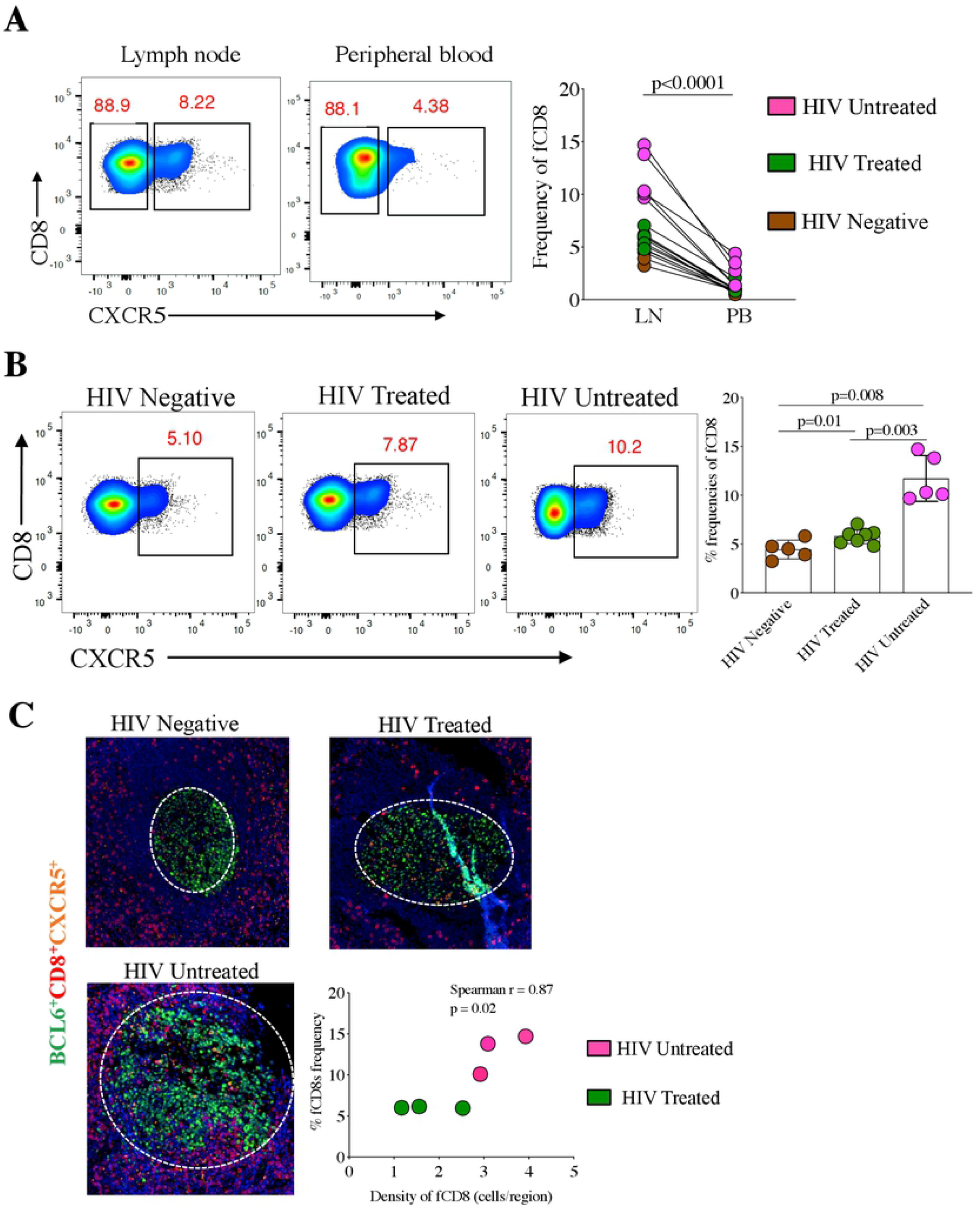
Phenotypic characterization of fCD8s in LN in HIV-1 infection. **(A)** Paired comparative analysis of the % frequency of fCD8s (CD8^+^CXCR5^+^) in lymph node (LN) and peripheral blood (PB) of 17 participants comprises of 5 untreated, 7 treated and 5 HIV negative individuals. Analysis shows a significantly magnitude of fCD8s in LN compared to PB. **(B)** Comparative analysis of HIV treated and untreated groups with HIV negative group showing a significant increase in fCD8s in HIV infected groups. **(C)** LN imaging showing the density of fCD8s within the germinal centre (GC). Correlation analysis showing significant positive correlation between the frequency of fCD8s measured by flow cytometry with the density of fCD8s in GCs quantified by imaging of fixed tissue (TissueQuest) in HIV treated and untreated groups.

We then evaluated the effect of HIV infection and viral antigen persistence on the induction of fCD8s in LNs, comparing participants in the FRESH cohort who were uninfected with subjects who were ART suppressed, as well as untreated donors. We observed a significantly higher frequency of fCD8s as a percentage of total CD8^+^ T cells in treated (p=0.01) and untreated donors (p=0.008) compared to uninfected donors **(Fig. 1B)**, and that ART limited the development of this phenotype (p=0.003) **(Fig. 1B)**. These data is consistent with previous studies that suggest persistent viral infection (16, 22) and/or inflammation (16) in ART-suppressed individuals drives the differentiation of fCD8s during HIV infection.

We next assessed if increased fCD8s in HIV treated and untreated individuals as compared to HIV negative individuals correlated with their localization in GCs using multicolour immunofluorescence microscopy and TissueQuest image analysis software. This technique allows simultaneous quantitative assessment of cellular phenotype and cell localization in tissues (30). We defined fCD8 as CXCR5^+^CD8^+^ T cells. Active GCs were identified by BCL6^+^ staining within B cell follicles. Image analysis readily revealed fCD8s localized in the GCs in HIV infected persons, in contrast to the lack of GC fCD8s in HIV negative persons **(Fig. 1C)**. Notably, we observed a significant positive correlation between the density of fCD8s localized in GCs and the frequency of fCD8s measured by flow cytometry in treated and untreated HIV infection (r=0.87, p=0.02) **(Fig. 1C)**, consistent with the notion that, viral infection stimulate proliferation of fCD8, which preferentially localize in GCs.

### Transcriptional and epigenetic factors are differentially expressed between human fCD8s and GCTfh

fCD8s are associated with HIV and tumour control (16, 18), but their differentiation conditions in humans are not known. Recent animal studies have defined the regulatory networks that govern the expression of CXCR5 in CD8^+^ T cells (16, 17, 22, 23). However, it is not clear if similar regulatory networks regulate CXCR5 expression in human CD8^+^ T cells. To address this question, we performed bulk RNA-Seq on FACS-sorted cells from the excised LNs of five HIV infected individuals (**supplementary Fig 1A**). Five separate cell populations were FACS-sorted from each individual: bulk fCD8s (CD3^+^CD8^+^CD45RA^-^CXCR5^+^), non-fCD8s (CD3^+^CD8^+^CD45RA^-^CXCR5^-^), naïve CD8^+^ T cells (CD3^+^CD8^+^CD45RA^+^CCR7^+^), GCTfh (CD3^+^CD4^+^CXCR5^high^PD1^high^) and non-Tfh (CD3^+^CD4^+^CXCR5^-^PD1^-^) (**supplementary Fig. 1B**). GCTfh, which constitutively express high levels of CXCR5, and naïve CD8^+^ T cells, which do not express CXCR5, served as positive and negative controls, respectively. Non-Tfh was included as additional control to compare with GCTfh. Principal component analysis (PCA) of 5 biological replicates separated all experimental groups in two dimensional space based on quantification of mRNA transcripts **(Fig. 2A)**. Despite minimal separation of fCD8s and non-fCD8s, there were 607 genes (FDR<0.1) that were differentially expressed between these two subsets (**supplementary data file**).

**Figure 2:**
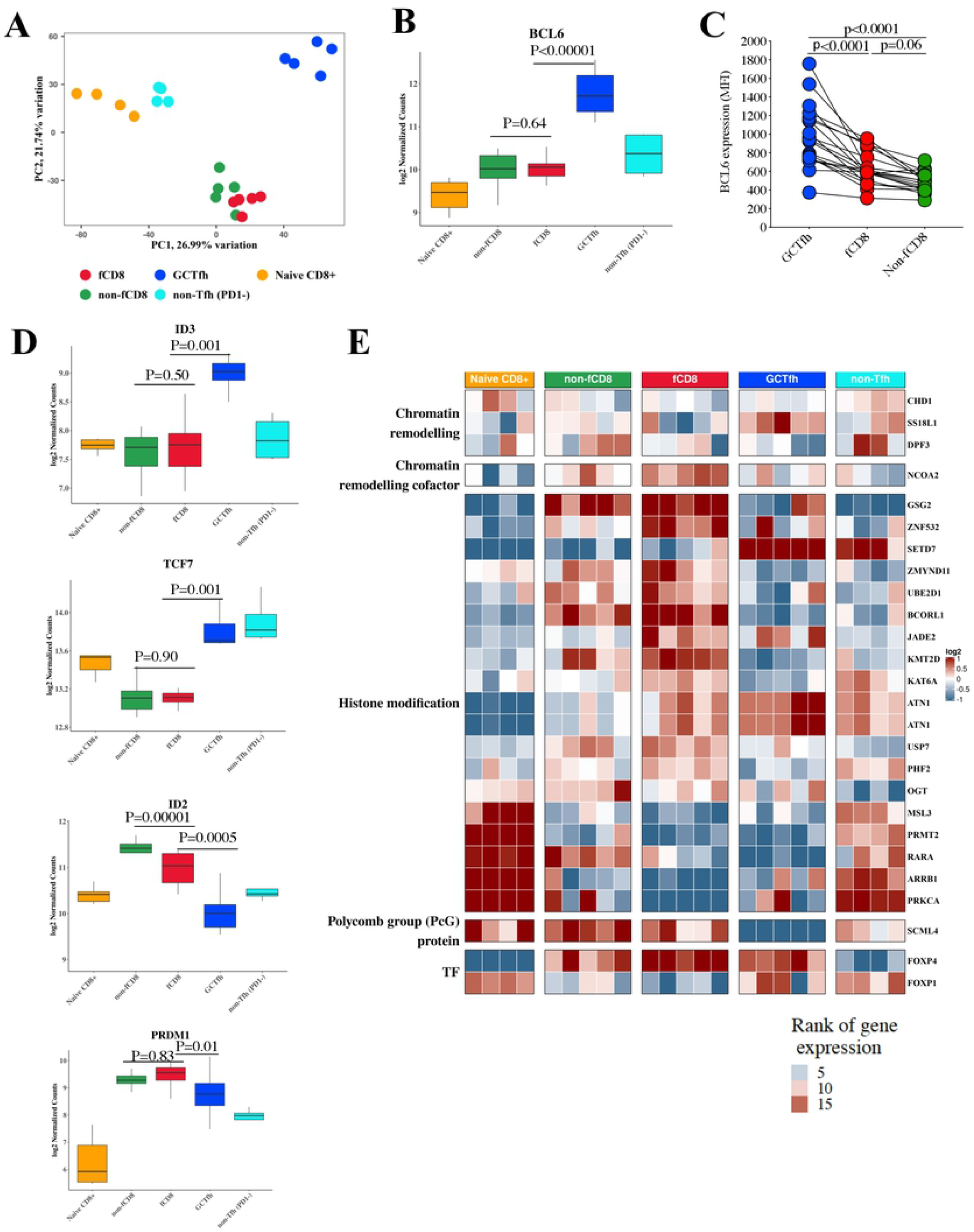
Lower expression of *BCL6* in fCD8s compared to GCTfh. **(A)** Principal component analysis of the RNA-Seq data from the four cell subsets, colour labelled according to cell subset. The top 500 genes by variance were used to construct the PCA plot. Clear separations are observed between the subsets with the fCD8s and non-fCD8s subsets showing closest proximity. **(B)** Statistical analysis showing significant greater magnitude *BCL6* expression in GCTfh compared to fCD8s and no difference between fCD8s and non-fCD8s. **(C)** Statistical analysis showing significant increase of BCL6 mean fluorescence intensity (MFI) in GCTfh compared with fCD8s and non-fCD8s. **(D)** Expression values of CXCR5 regulating genes. Batch and patient corrected transcripts per million (TPM) values for selected genes previously shown to be involved in the regulation of CXCR5 expression. FDR values are obtained from the differential expression analysis using the sleuth package in the R statistical environment. **(E)** Ranked expression of selected epigenetic modifiers. Epigenetic modifiers were grouped according to functional attributes, i.e. chromatin remodeling, histone chaperone, histone modification and by transcription activity. Genes were ranked from highest (red) to lowest (blue) expression. Each column represents the expression level for a particular patient as labelled on the x-axis.

We first analysed genes that have previously been implicated in *CXCR5* regulatory circuitry in animal studies, beginning with BCL6, which has been described as the master regulator of *CXCR5* gene expression in GCTfh and murine fCD8s (16, 22, 32, 33). We found that BCL6 was highly expressed in GCTfh relative to fCD8s (p<0.00001) and non-Tfh (p=0.16) **(Fig. 2B)**. Notably, there was no difference in BCL6 expression between fCD8s and non-fCD8s (p=0.64) **(Fig. 2B)**, contrary to murine studies (16, 22). To determine if BCL6 expression levels correlate with protein levels, we measured BCL6 expression by flow cytometry. Consistent with the transcriptional analysis, BCL6 expression was significantly lower in fCD8s compared to GCTfh (p<0.0001) **(Fig. 2C)**. Together these data indicate a fundamental difference in transcriptional circuitry that regulate CXCR5 expression in follicular CD4^+^ relative to CD8^+^ T cells. The data also suggest that BCL6 may not be a critical regulator of CXCR5 expression in human CD8^+^ T cells.

Next, we investigated other genes that were similarly expressed between fCD8s and GCTfh in mice, and were reported to be part of the CXCR5 transcription circuitry. They include: *Id3, Id2, TCF7* (gene coding for *TCF-1*) and *PRDM1* (22). Again, contrary to what was reported LCMV mouse models (16, 22), Id3 and TCF7 were significantly downregulated in human fCD8s compared to GCTfh (Id3: p<0.0001, TCF-1: p<0.0001), with no apparent difference between fCD8s and non-fCD8s (Id3: p=0.50, TCF-1: p=0.90) (**Fig 2D**). Similarly, *Id2*, which is a negative regulator of CXCR5 expression was significantly higher in fCD8s compared to GCTfh (Id2: p=0.0005). *PRDM1* that has been shown to antagonize GCTfh differentiation (32), was significantly higher in fCD8s compared to GCTfh (p=0.01) **(Fig. 2D)**. Notably, *Id2* was significantly expressed between fCD8 and non-fCD8 but not *PRDM1* (Id2: p=0.00001, PRDM1: p=0.83). Together, these data suggest that the common transcriptional regulators of CXCR5 expression in GCTfh and fCD8, described in murine studies, is true for human GCTfh but not fCD8s. These data suggest that an alternative transcription circuitry may be involved in regulating CXCR5 expression in human CD8^+^ T cells.

To gain further insight into the transcriptional mechanisms responsible for the *CXCR5* gene regulation in human CD8^+^ T cells, we focused on the genes found to be differentially expressed between lymphoid fCD8s and non-fCD8s by RNA-Seq analysis. We identified 43 genes (≈7% of differentially expressed genes, FDR<0.1) that encode factors regulating epigenetic processes (epigenetic factors) such as chromatin remodelling, histone modification and DNA methylation (34) **(Fig. 2E and extended data in supplementary Fig. 2)**. These data provided the first hint that specific epigenetic mechanisms maybe directly involved in regulating CXCR5 in human CD8^+^ T cells.

### The *CXCR5* gene locus is tightly regulated by DNA methylation and chromatin landscape in human lymphoid CD8^+^ T cells

Epigenetic regulators were among the most highly differentially expressed genes between fCD8s and non-fCD8s; thus, we hypothesized that distinct epigenetic mechanisms, such as changes to DNA methylation and/or chromatin landscape, regulate the expression of CXCR5 in human CD8^+^ T cells. To obtain experimental evidence to address this, we first measured DNA methylation levels proximal to *CXCR5* from the same cell populations used for RNA-Seq, using loci-specific bisulfite-treated DNA sequencing. We FACS-sorted GCTfh, fCD8s, non-fCD8s, and naïve CD8^+^ T cells from LNs. We did not include non-Tfh in this experiment due to sample availability and that we could only FACS-sort 4 subsets at a time. We extracted DNA from 3 biological replicates for sequencing. DNA methylation levels were measured in CpG islands within 300 bp upstream to 200 bp downstream of the CXCR5 TSS. We observed significantly higher methylation levels proximal to the CXCR5 promoter region in naïve CD8^+^ T cells (average methylation 88%), non-fCD8s (average methylation 69%). In contrast, fCD8s (average methylation 7%) and GCTfh (average methylation 6%) had minimal levels of methylation at equivalent sites **(Fig. 3A and B)**.

**Figure 3:**
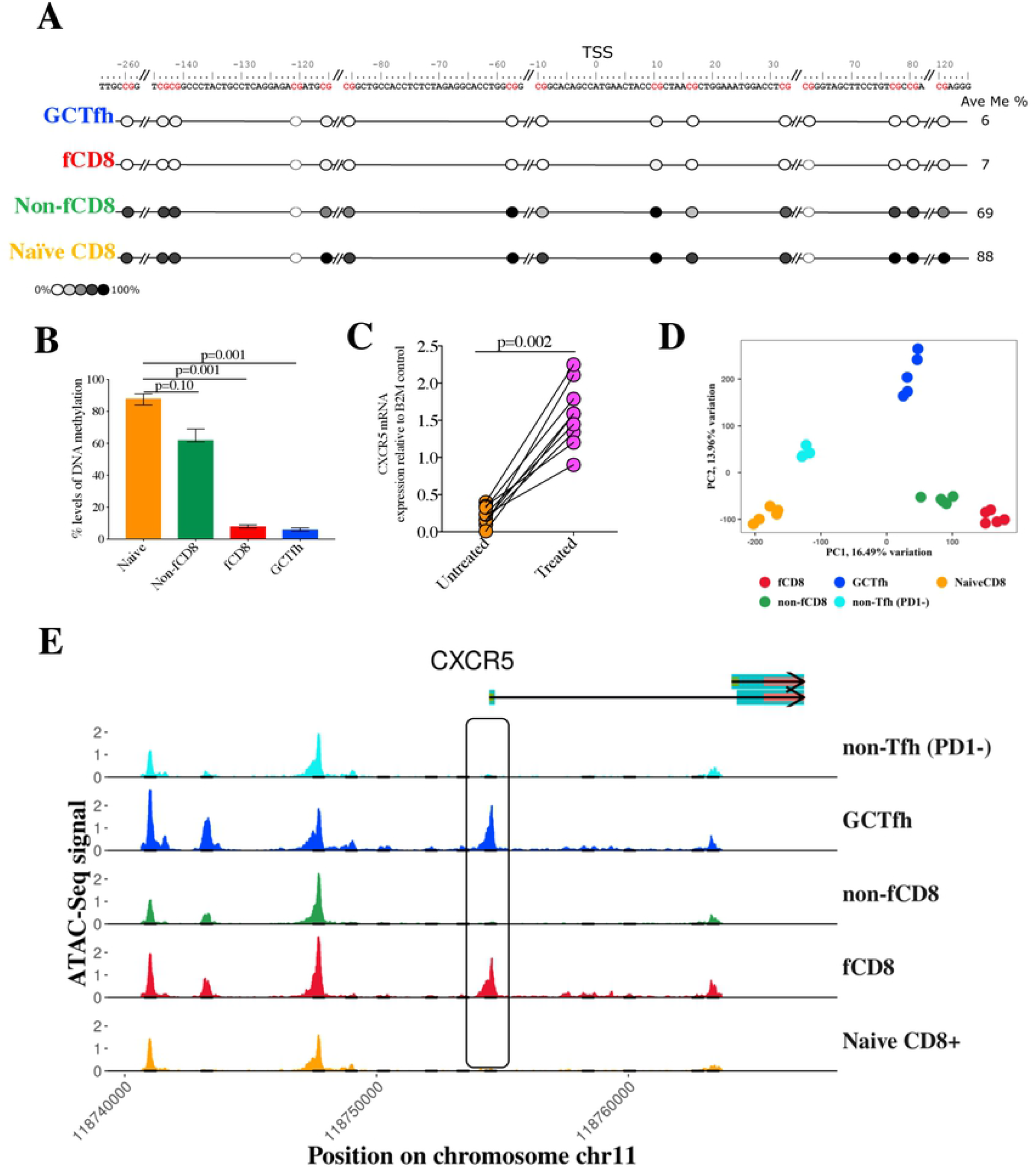
Epigenetic regulation of CXCR5 expression. **(A)** Quantitative measurement of DNA methylation levels within specific cell subsets; GCTfh, fCD8s, Non-fCD8s and Naïve-CD8^+^ T cells were determined using the EpiTYPER^®^ DNA Methylation Analysis. Methylation levels were measured from bi-sulfite treated genomic DNA, followed by PCR amplification of a 500bp fragment containing 15 CpG sites (red letters). The naïve- and non-fCD8s cells show higher levels of methylation within several sites (darker circles), while the GCTfh and fCD8s show lower levels of methylation (lighter circles), suggesting DNA methylation interference with *CXCR5* gene transcription. The position of CpG sites are represented relative to the transcription start site (TSS). **(B)** Percentage levels of methylation are depicted in bar graph for each subset analyzed across the 15 CpG sites. **(C)** Non-fCD8s were FAC-sorted and treated for 24 hours with 10μM 5-aza-2’-deoxycytidine (Aza drug), a DNA methyltransferase inhibitor that causes hypomethylation of DNA. Fold change relative to the B2M house keeping control indicated significant increase in the CXCR5 expression levels after treatment, indicating DNA methylation as potential mechanism limiting transcription of *CXCR5* gene. **(D)** PCA plots obtained from the ATAC-Seq cut count data. The top 10% of ATAC-Seq peaks (merged between subsets) by variance were used to create the PCA plot. **(E)** Overview of the ATAC-Seq signal around the *CXCR5* gene loci. ATAC-Seq signal is shown for different marked (in grey) loci where differential binding was detected in at least one sample. The black box shows the TSS region where there is clear equivalence between fCD8s and GCTfh ATAC-Seq signals, while very low signal was observed for both non-fCD8s and naive CD8^+^ T cells.

To determine if methylation was responsible for *CXCR5* gene silencing, we incubated FACS-sorted non-fCD8s with 10μM of 5’-aza-2-deoxycytidine (Aza), which inhibits the enzymatic activity of DNA methyl transferases (38). After 24 hours of incubation, we measured CXCR5 mRNA transcript levels by digital droplet PCR (ddPCR). We found that Aza treatment significantly increased CXCR5 mRNA levels (p=0.002) **(Fig. 3C)**. Together, these data suggest that *CXCR5* gene locus-specific DNA methylation is involved in repressing the *CXCR5* gene in human non-fCD8s.

In addition to showing that DNA methylation is likely involved in repressing CXCR5 transcription, the RNA-Seq dataset revealed several other differentially expressed genes involved in epigenetic regulatory processes such as chromatin remodelling and histone modification. Thus, to gain comprehensive mechanistic insights into the epigenetic processes that regulate *CXCR5* gene expression in CD8^+^ T cells, we used the Assay for Transposable-Accessible Chromatin using Sequencing (ATAC-Seq). This technology identifies genome wide accessible regions and can be used to identify transcription factor (TF) footprinting and nucleosomal positioning, all of which cooperatively regulate gene expression (39, 40). Briefly, ATAC-Seq analysis was performed on the DNA samples isolated from the same lymphoid cell populations used for RNA-Seq studies (**supplementary Fig. 1B)**. We performed a PCA on the top 10% variably accessible regions, revealing clear delineation of cell subsets based on the chromatin accessibility profiles **(Fig. 3D and supplementary Fig. 3A)**. We calculated a set of 66,514 open chromatin regions (OCRs) that appeared in at least one of the subsets. The subset separation was strikingly similar to the PCA plot for RNA-Seq data (**see Fig. 2A**), revealing significant overlap between accessibility and gene expression. Indeed, there was a strong association between chromatin accessibility and gene expression between fCD8s and non-fCD8s (R^2^= 0.54) **(supplementary Fig. 3B)**.

Next, we profiled accessibility of the *CXCR5* gene, revealing a closed chromatin conformation at the TSS of the *CXCR5* gene in non-fCD8s, naïve CD8^+^ T cells and non-Tfh. In contrast, fCD8s and GCTfh had open chromatin conformation at the equivalent site **(Fig. 3E)**. These data confirm that chromatin accessibility also contributes to the repressed state of the *CXCR5* gene in non-fCD8s and naïve CD8^+^ T cells. The observed DNA methylation and closed chromatin structure of the CXCR5 TSS are consistent with the notion that DNA methylation promotes nucleation of repressed chromatin structure encompassing the *CXCR5* gene region (27, 41).

To identify epigenetic factors that may directly regulate chromatin accessibility of the *CXCR5* gene, we next performed a TF binding motif search around the *CXCR5* TSS. We restricted the motif search to regions that were inputted to have TF footprints proximal to the TSS (42, 43). Our analysis revealed that fCD8s and GCTfh shared binding motifs at the *CXCR5* gene TSS for several epigenetic regulatory proteins, namely Pit-Oct-Unc (POU) family: POU2F3, POU3F1, POU3F3, E2F6, and ZNF384 (**Fig. 4A**). Given that POUs-TFs function as pioneer factors that interact with the closed chromatin at enhancer and/or promoter regions to open up regions for transcriptional activities (25, 44–46), and the fact that POU2F3, POU3F1 and POU3F3 binding sites were observed for both fCD8s and GCTfh, these data suggest that these three pioneer factors may be directly involved in opening the chromatin structure at the CXCR5 TSS.

**Figure 4:**
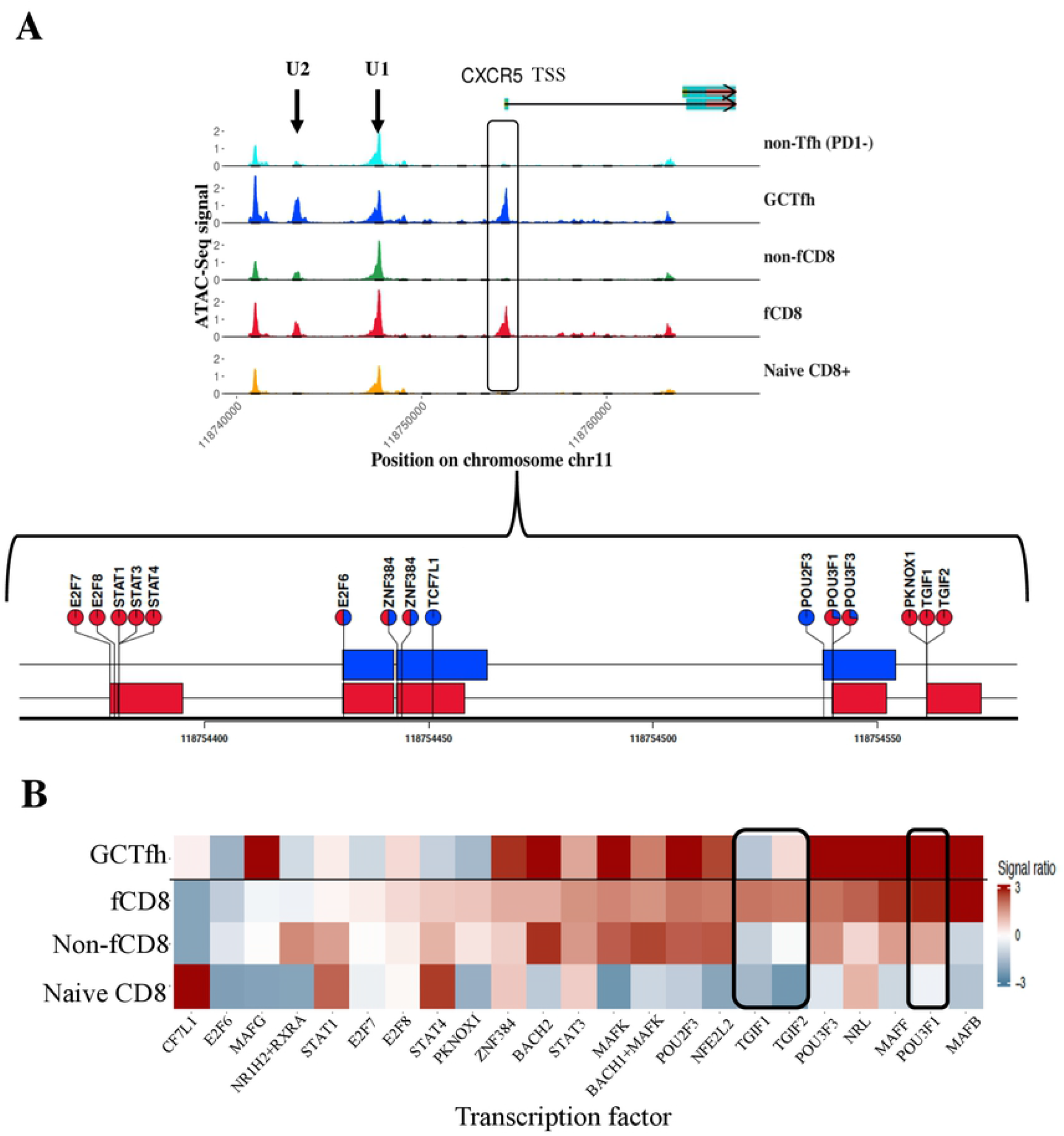
Shared and unique transcriptional factor footprint proximal to the *CXCR5 gene*. **(A)** Footprints in selected regions predicted footprinted regions respective cell subsets. The pie charts show the relative Wellington bootstrap scores for each subset against all others acting as a proxy for the relative TF activity observed in that region. The bars indicate the extent of the predicted TF footprint, with colours assigned to each subset. Footprints with unassigned TFs are also included. **(B)** Assignment of TF to subsets. Enrichment of TF motifs (restricted differential imputed footprints between subsets) of each subset is depicted in the heatmap. The TFs are sorted in ascending order of importance through the signal ratio in the fCD8 subset.

We next looked for TF binding sites upstream of the *CXCR5* TSS. ATAC-seq analysis identified two peaks upstream of the *CXCR5* TSS, likely representing enhancer regions which we labelled U1 (−6.5kb), and U2 (−11kb) (**supplementary Fig. 4A)**. We performed a TF motif search within these regions for each subset to identify specific TFs that bind in this region and found that MAF was highly enriched in fCD8s and GCTfh, while TGIF1 and TGIF2 were enriched in fCD8s but not in GCTfh (**Fig. 4B**). Together, these data suggest that POU epigenetic pioneering factors mediate the opening of chromatin around the *CXCR5* TSS and that MAF, TGIF1 and TGIF2 are key TFs in CXCR5 expression in human CD8^+^ T cells.

### Weighted Gene Correlated Network Analysis (WGCNA) reveals alternative pathway involved in the expression of CXCR5 in human CD8^+^ T cells

Cell differentiation involves complex interplay between transcription factors that progressively dictate their phenotype and function. To identify molecular circuitry that regulate *CXCR5* gene expression in human CD8^+^ T cells, we performed Weighted Gene Correlated Network Analysis (WGCNA) on the ATAC-Seq and RNA-Seq data sets. WGCNA is a network analysis that is used to identify modules of highly co-expressed genes using RNA-Seq data (47) or chromatin-accessible gene networks using ATAC-Seq data (48, 49). The program assigns a identifier to each module as an identification mark. We first applied this network analysis on the ATAC-Seq data to identify chromatin accessibility networks that cooperatively regulate CXCR5 gene accessibility. We hypothesized that the mechanisms governing chromatin accessibility may not act on open chromatin regions (OCRs) in isolation, but rather are grouped into programs that change the accessibility of multiple chromatin loci.

We performed WGCNA on 12,000 ATAC-Seq peaks after excluding sites with high technical variance and retaining regions proximal to genes that were differentially expressed. Interestingly, we observed that the *CXCR5* TSS and U2 OCRs were both assigned by WGCNA to module 5 (**Fig. 5A**). These data suggest that the U2 (enhancer region) interacts with the TSS to promote CXCR5 transcription. Notably, enrichment analysis on module 5 revealed striking similarity in accessibility pattern between fCD8s and GCTfh (**Fig. 5B**), despite the clear difference in overall genome-wide accessibility between the two cell subsets as depicted in ATAC-Seq PCA plot (**Fig. 3D**). Importantly, these data identify regions that regulate *CXCR5* gene accessibility that are shared between fCD8s and GCTfh.

**Figure 5:**
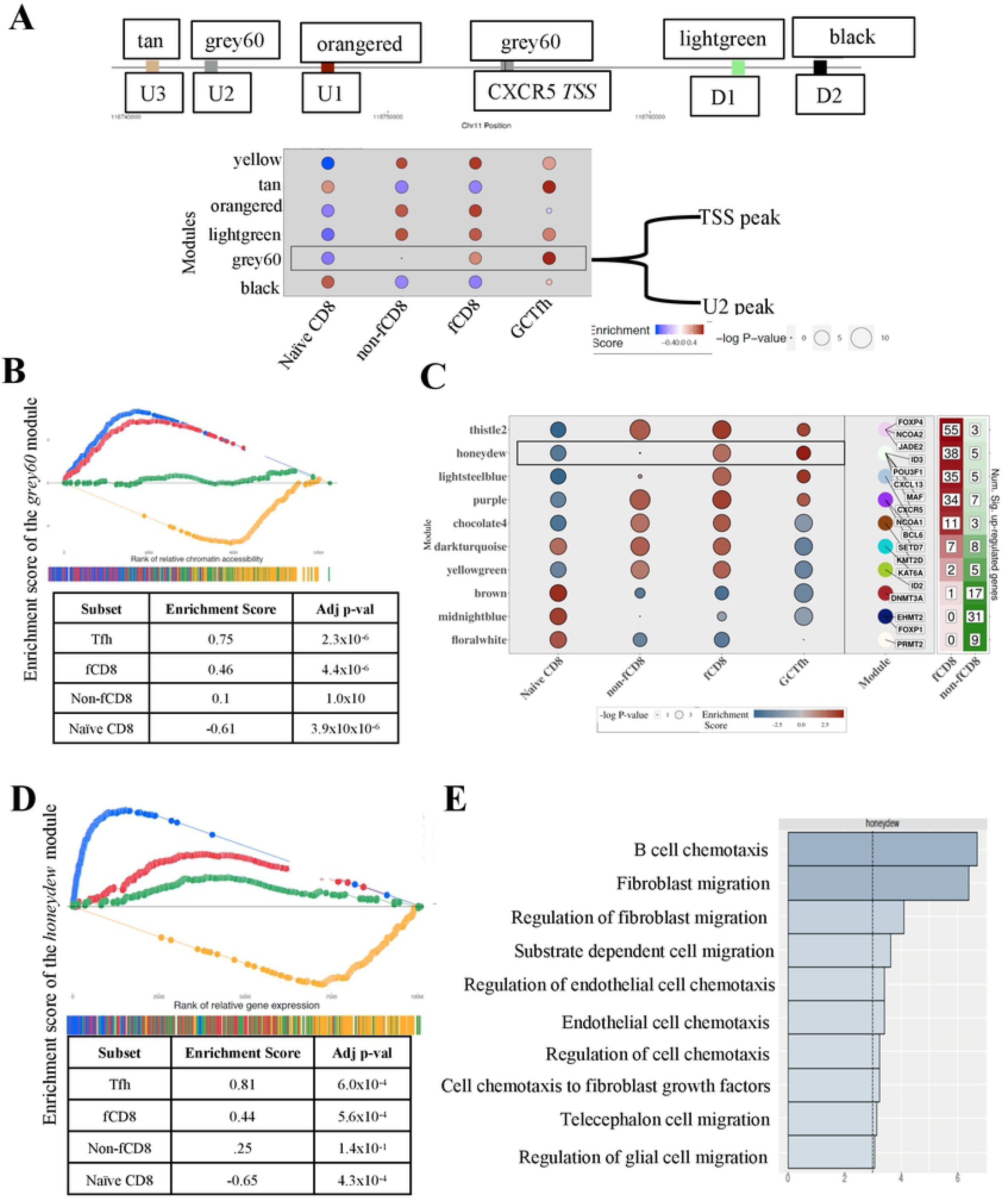
Regulatory pathways influencing CXCR5 expression. **(A)** The top figure shows the OCR regions observed in at least one of the cell subsets. Peaks are either prefixed with U to indicate upstream, or D to indicate downstream of the CXCR5 TSS. The colours represent the WGCNA modules. Module names appear at peak regions. ATAC-Seq WGCNA around the *CXCR5* gene region. Modules are sized according to enrichment and significance. The highlighted module 5 contains both the TSS of *CXCR5* and the U2 region. **(B)** Gene set enrichment analysis (GSEA) plot of the module 5 of ATAC-Seq WGCNA enrichment values. Peaks belonging to the module 5 for each subset are plotted according to the rank within each subset. High correspondence and enrichment are seen for the GCTfh and fCD8s subsets, while no enrichment is shown for non-fCD8s and negative enrichment is shown for naïve CD8^+^ T cells. **(C)** Overview of the RNA-Seq WGNCA modules. Selected modules are shown. The modules are named according to their GSEA score. Positive values indicate positive enrichment. The size of the module corresponds to the −log P-value. The panel to the right indicates the number of genes that are up-regulated in fCD8 and non-fCD8 for each module. **(D)** GSEA analysis shows the overall enrichment of the CXCR5 containing in module 2, with corresponding enrichment scores and significance values. The bottom bar shows the concentration of genes within a subset according to the rank of expression. **(E)** GO enrichment of the module 2 showing positive enriched GO terms in the module 2 ranked according to significance. Cell migration is an important factor in the module 2.

Having identified gene accessible regions that are potentially involved in CXCR5 accessibility in both fCD8s and GCTfh at DNA level, we next performed WGCNA analysis on the RNA-Seq data to define CXCR5 gene regulation at mRNA level. We constructed WGCNA networks using the four cell subsets and 20,987 genes sufficiently expressed in fCD8s and non-fCD8s. The resultant network consisted of 91 modules, each containing a set of highly co-expressed genes. We batch normalized the data to account for heterogeneity of expression between participants and used the expression values to calculate gene set enrichment analysis (GSEA) for each subset for the 91 detected modules. We observed a significant enrichment of *CXCR5, MAF, Id3, POU3F1* and *CXCL13* genes in module 2, which was shared by fCD8s and GCTfh **(Fig. 5C)**. Importantly, ATAC-Seq data identified motifs from footprints for the same set of genes as ATAC-Seq data in U2 and TSS regions of the *CXCR5* gene **(Fig. 4B; supplementary Fig. 4A**). Notably, GSEA demonstrated significant enrichment of GCTfh and fCD8s subsets in module 2 **(Fig. 5D)**. Gene ontology (GO) analysis on the CXCR5-centric module 2 showed enrichment of terms associated with “cell migration” **(Fig. 5E)**, suggesting that a subset of genes governing the expression of CXCR5 in human CD8^+^ T cells are intricately involved in cell migration. Collectively, our data identify *MAF, Id3* and *POU3F1* as key genes involved in driving the expression of CXCR5 in human CD8^+^ T cells.

Based on the experimental and computational data presented in this study, we propose the following model for the expression of CXCR5 on human CD8^+^ T cells in lymphoid tissues; T cell receptor (TCR) stimulation of lymphoid tissue naïve CD8^+^ T cells leads to a stepwise chromatin plasticity driven by pioneering factor (POU3F1) and DNA demethylation that cooperatively open up chromatin at the CXCR5 TSS and promoter region. Chromatin relaxation allows the recruitment of transcriptional machinery including *MAF, Id3, TGIF1, TGIF2 and CXCL13* that drive the expression of CXCR5 (see details of the proposed model in **supplementary Fig. 5**).

### Low CXCR5 expression on fCD8s impacts their migratory capacity to the germinal centers

We next investigated the observed lower expression of CXCR5 in fCD8s relative to GCTfh, which is thought to attenuate their migratory capacity into B cell follicles (20, 50). We first compared CXCR5 protein expression levels and found significantly higher expression in GCTfh compared to fCD8s (p=0.0001) **(Fig. 6A)**, consistent with previous reports (29). This was true for mRNA levels as well **(Fig. 6B)**. We then performed a trans-well experiment to assess if expression of CXCR5 affects the rate of fCD8s chemotaxis towards a CXCL13 gradient. Indeed, fCD8s exhibited significantly lower chemotaxis capacity compared to GCTfh (p=0.0001) **(Fig. 6C)**. Moreover, a GO analysis on the RNA-Seq data showed enrichment of genes associated with cell migration/leukocyte migration in fCD8s relative to non-fCD8s **(Fig. 6D)**. Together, these data confirm that lower expression of CXCR5 reduces chemotaxis capacity of fCD8s towards CXCL13.

**Figure 6:**
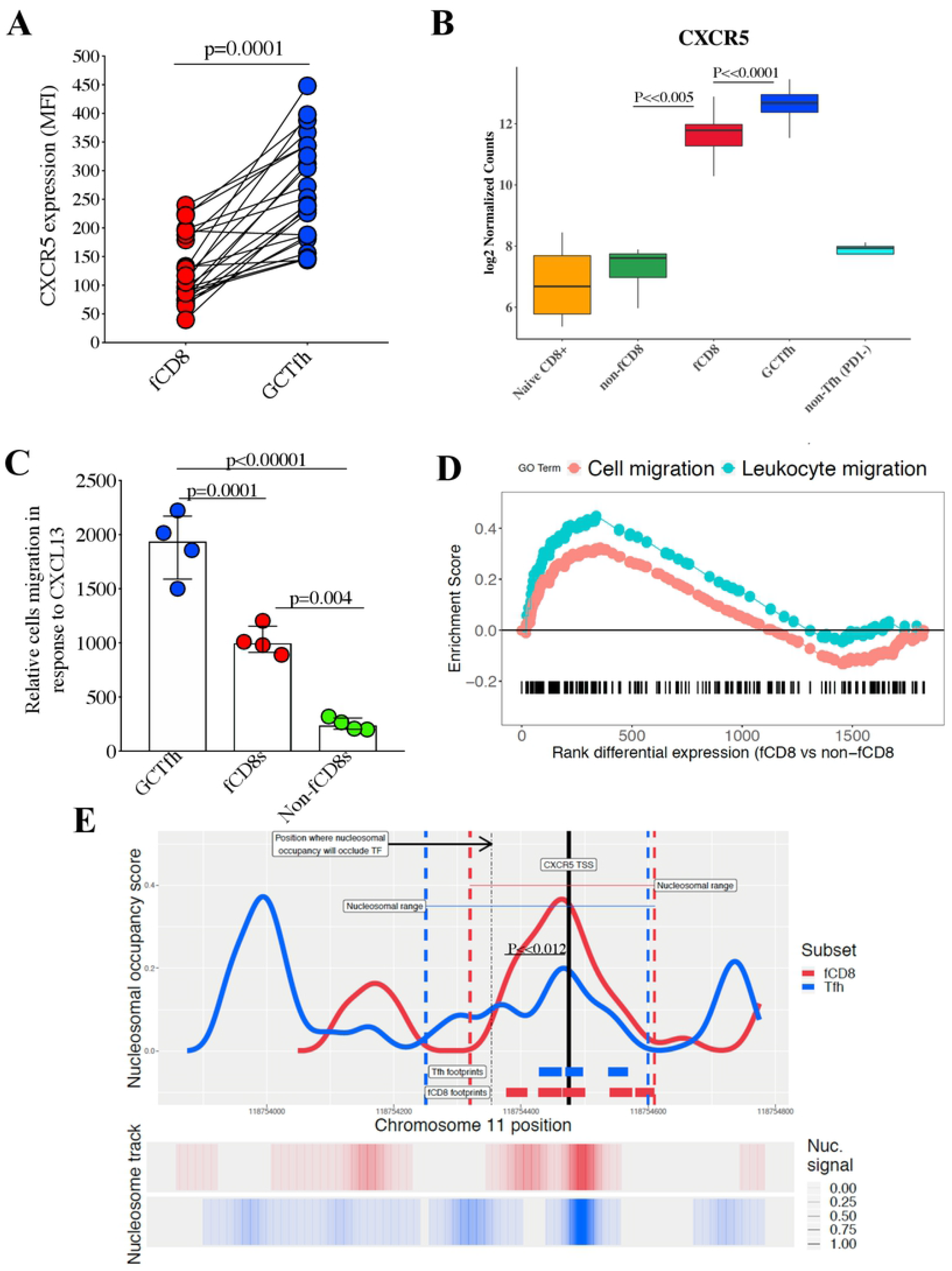
CXCR5 expression level on fCD8s impacts their migration to the germinal centers. **(A)** Mean fluorescence intensity (MFI) of CXCR5 on fCD8s and GCTfh shows significant increase in the expression of CXCR5 on GCTfh compared to fCD8s. **(B)** RNA-Seq expression values of CXCR5 showing the batch-normalized expression values in different cell subsets. **(C)** Relative migration of GCTfh, fCD8s and non-fCD8s subsets in response to CXCL13; a ligand for CXCR5. Graph shows the number of cells that migrated in each subset after 3 hours **(D)** The GSEA plot of GO terms between fCD8s and non-fCD8s Cell migration and Leukocyte migration shows the ranked differential expression of genes belonging to these terms between the fCD8s and non-fCD8s subsets. **(E)** The figure depicts the nucleosomal occupancy scores (top line plot) and the nucleosomal signal (bottom heatmap) as produced by NucleoATAC around the TSS region of CXCR5 in fCD8 (red) and GCTfh (blue) subsets. The colored vertical dashed lines show the range of predicted nucleosomal occupancy. The thin dashed line shows the approximate location of the nucleosomal dyad where the nucleosome will occlude the TSS region. Height of the occupancy score shows the fraction of nucleosomal sized fragments at the chromosome 11 position. Predicted transcription factor footprints are shown as bars for the respective cell subsets. The heatmap shows the calculated nucleosomal signal from the ATAC-Seq data and shows a higher degree of nucleosomal translocation in the 5’ direction in GCTfh compared to fCD8s.

### Reduced turnover rate of the nucleosome at the promoter region of *CXCR5*

Next, we investigated the molecular basis of low CXCR5 expression on fCD8s. Given high frequency of methylated CpG islands in the *CXCR5* gene, which tend to attract nucleosomes (27, 51), we evaluated nucleosomal occupancy at the TSS. We hypothesized that nucleosome positioning and occupancy around the TSS would interfere with the transcriptional machinery resulting in mitigated gene expression (52). To test this, we used the NucleoATAC tool (53) to impute the presence of nucleosomes in and around the *CXCR5* gene. Interestingly, the presence of nucleosomes was imputed in both fCD8s and GCTfh at the TSS. However, nucleoATAC revealed higher nucleosomal occupancy in predicted TF footprint regions around the TSS in fCD8s, whereas GCTfh exhibited less nucleosomal occupancy in the same region **(Fig. 6E)**. Computationally, the nucleosomal occupancy was calculated for a wider range upstream of the TSS in GCTfh than fCD8s (blue and red dotted lines), which extended beyond the point where a nucleosome may occupy TF binding regions (black dashed line) **(Fig. 6E)**, suggesting that positioning of nucleosome at the TSS may interfere with optimal transcription of *CXCR5* in fCD8. Given that nucleosome occupancy results in the enrichment of reads with longer insert sizes in the ATAC-Seq data, typically greater than 147 bp. We used this knowledge as a proxy for nucleosomal occupancy and performed a Fisher’s exact test to compare longer to shorter read ratio over the imputed dyad of the TSS nucleosome (53). We found that the ratio of longer to shorter reads were 2.38 times what was calculated for GCTfh (p=0.012). Collectively, these data suggest higher nucleosomal occupancy in fCD8s compared to GCTfh at the TSS region. Computational simulation of nucleosomal occupancy confirmed the notion that nucleosomal occupancy interferes with transcriptional machinery, reducing the transcription of the *CXCR5* gene in fCD8s **(supplementary Fig. 6A and B)**.

## Discussion

Understanding regulation of CD8^+^ T cell trafficking to B cell follicles has far reaching implications for developing strategies to eradicate HIV infected cells in B cell follicles and to treat B cell derived malignancies. This study set out to address two key questions. First, we investigated why the majority of CD8^+^ T cells that reside in LNs do not express CXCR5, the chemokine receptor required for cellular trafficking into LN follicular areas. Second, we interrogated the molecular mechanisms that regulate differential levels of CXCR5 expression on fCD8s relative to GCTfh.

To answer the first question, we studied two antigen-experienced CD8^+^ T cell subsets termed fCD8s and non-fCD8s that were phenotypically matched except for the expression of CXCR5 on the cell surface. Locus-specific bisulfite-treated sequencing and genome-wide chromatin accessibility data identified DNA-hypermethylation and closed chromatin structure as two epigenetic mechanisms that are involved in repressing CXCR5 expression in human non-fCD8s.

For the second question, we focused the analysis on CXCR5^+^ subsets, fCD8s and GCTfh, because of the significant difference in the levels of CXCR5 expression and trafficking kinetics between the two subsets. We showed that fCD8s had reduced CXCR5 expression compared to GCTfh, and were less efficient at trafficking towards CXCL13 chemokine. Importantly, we identified marked differences in nucleosomal occupancy and positioning between these two subsets, suggesting a plausible mechanism moderating the expression of CXCR5 in fCD8s. Taken together, our data show that CXCR5 expression in CD8^+^ T cells is tightly controlled by at least three key epigenetic mechanisms: DNA methylation, chromatin structure and nucleosomal occupancy.

Conceptualization of this study was motivated by three studies in mice that recently described a subset of CXCR5 expressing CD8^+^ T cells termed fCD8s because of their ability to accumulate in B cells follicles (16, 17, 22). Strikingly, the murine models showed that the transcriptional profile of fCD8s looks similar to that of GCTfh but not non-fCD8s. More importantly, the murine studies showed that following lymphocytic choriomeningitis virus (LCMV) infection, fCD8s readily accumulated in B cell follicles and were able to eradicate infected GCTfh (16, 17, 22). A subsequent rhesus macaque study showed similar results (23). Thus, we asked if fCD8s were also increased in HIV infection in human LNs and if their differentiation profile was similar to that described in mice. Indeed, our data show increased frequency of fCD8s in LN of HIV infected individuals compared to uninfected individuals. Initiation of antiviral therapy mitigated the fCD8s differentiation, suggesting that fCD8s induction is antigen driven, as described in animal studies (20). Increase fCD8s is probably not unique to HIV-1 infection but rather a more generalized immune response to viral infection in LNs.

Given that murine studies identified several TFs that were common between fCD8s and GCTfh, including BCL6, Id3, Id2, PRDM1 and TCF-1 (16, 17, 22, 32, 54), we investigated whether similar TFs were operating in human LN CD8^+^ T cells in the setting of HIV infection. We sorted LN fCD8s and non-fCD8s from HIV infected individuals using sorting panels based on similar markers used in murine studies. RNA-Seq analysis showed that TF expression profiles in human GCTfh cells were similar to those reported in mice (22). In contrast to the murine studies, we found significant differences in TF expression profiles between human GCTfh and fCD8s. In fact, our data show that TF expression profiles in human fCD8s were more similar to non-fCD8s than GCTfh. This indicates that most of the TFs that are critical for fCD8 differentiation in mice might not be essential for human fCD8 differentiation. Taken together, these results suggest that other mechanisms may regulate CXCR5 expression in human CD8^+^ T cells.

Our RNA-Seq data indicate that epigenetic mechanisms play a major role in *CXCR5* gene regulation. Locus-specific bisulfite-treated sequencing revealed hypermethylation in CpG islands proximal to promoter regions of subsets that lack CXCR5 expression (non-fCD8s and naïve CD8^+^ T cells) and reduced methylation levels in CXCR5 positive cells (fCD8s and GCTfh). Moreover, inhibition of enzymatic activity of methyltransferase using aza treatment increased CXCR5 expression in CXCR5 negative cells, thus, providing compelling evidence that DNA methylation is a major epigenetic mechanism involved in silencing CXCR5 expression (36). ATAC-Seq data revealed closed chromatin conformation at the CXCR5 TSS in non-fCD8s. It is a well-known phenomenon that DNA methylation increases nucleosome compaction and rigidity (41), therefore, greater DNA methylation is the probable cause of the observed condensed chromatin at the CXCR5 TSS and the corollary silencing of the *CXCR5* gene in non-fCD8s.

To identify key genes and pathways involved in *CXCR5* gene regulation, we performed WGCNA on the ATAC-Seq and RNA-Seq data. This allowed us to identify circuits of correlated chromatin accessibility as well as gene expression. WGCNA analysis identified modules of highly correlated open chromatin regions which indicates chromatin accessibility of the CXCR5 promoter region is part of a larger epigenetic circuit. We identified an important module that contains TSS and U2 peaks (a putative enhancer region). Strikingly, this module was highly enriched in both fCD8s and GCTfh which suggest that similar epigenetic circuitry shared between fCD8s and GCTfh in the context of regulation *CXCR5* gene accessibility. Furthermore, we used WGCNA to identify transcriptional modules that govern the expression of CXCR5 in human CD8^+^ T cells. From this analysis arose a module containing CXCR5, MAF, Id3, POU3F1 and CXCL13 which was enriched for fCD8s and GCTfh. GSEA on this module confirms a stepwise significance of genes skewed for GCTfh, followed by fCD8s, further implicating different regulatory pathway for CXCR5 expression in human CD8^+^ T cells. Indeed, GO analysis of this module clearly demonstrated chemotaxis and B cell migration as the key modules common to the two cell subsets. This implies that the transcriptional factors governing the expression of CXCR5 in human CD8^+^ T cells, such as Id3, MAF and POU3F1, were mostly contained in the same module.

Having identified the epigenetic processes that repress CXCR5 expression on non-fCD8s, we next focused on investigating molecular mechanisms that mitigate the expression of CXCR5 on fCD8s. Flow cytometry data and *in vitro* chemotaxis experiments suggest that lower expression level of CXCR5 in fCD8s contributes to the inefficient infiltration of B cell follicles observed in our imaging experiments. Importantly, we identified nucleosomal occupancy as a plausible molecular mechanism that likely lowers CXCR5 expression in fCD8s. We observed significant differential nucleosomal positioning at the TSS of fCD8s relative to GCTfh which suggests nucleosomal positioning as a plausible molecular mechanism.

A notable limitation of this study is that we could not profile the histone modification pattern around the *CXCR5* gene in fCD8s and non-fCD8s, due to insufficient sample availability for ChIP-Seq. Nonetheless, ChIP-Seq data in a B cell line that expresses higher levels of CXCR5 shows H3K4me2, which denotes open chromatin within the accessible regions around the *CXCR5* gene (GM12787 <u>(ENCODE Project Consortium 2012)</u>), consistent with our findings.

In conclusion, our data provide evidence of key epigenetic and transcriptional processes that intricately orchestrate the regulation of the *CXCR5* gene in human CD8^+^ T cells. Importantly, we identified a putative transcription circuitry that includes Id3, MAF and POU3F1, along with epigenetic mechanisms including DNA methylation, chromatin structure and nucleosomal occupancy as potential targets for inducing CXCR5 expression on human CD8^+^ T cells. Manipulation of these processes has the potential to enhance trafficking of CD8^+^ T cells to B cell follicles where they are needed to eradicate HIV infected cells or cancerous cells.

## Materials and methods

### Human samples

Fresh human inguinal lymph nodes (LNs) were obtained from participants enrolled at the Prince Memorial Mshiyeni Hospital, Umlazi township, Durban, South Africa. Demographic and clinical characteristics of the study participants are summarized in **Table 1**. A section of the excised LN was processed for tissue imaging and the remaining section was meshed to isolate lymph node mononuclear cells (LNMCs). LNs were homogenized using a syringe plunger and passed through a cell strainer (BD Biosciences Germany) to make a single-cell suspension. Mononuclear cells were isolated using RPMI medium (Sigma-Aldrich, St. Louis, MO) containing 10% heat-inactivated fetal calf serum (R10 medium). Extracted LNMCs were frozen for downstream experiments. All protocols were approved by the Biomedical Research Ethics Committee of the University of KwaZulu-Natal and the Massachusetts General Hospital Institutional Review Board.

### Flow cytometry and cell sorting

For phenotypic characterization, cells were surface stained with cell-viability dye (Fixable Blue dead cell stain kit, Invitrogen), followed by anti-CD3-BV711 (BioLegend), anti-CD4-BV650 (BD Biosciences), anti-CD8-BV786 (BD Biosciences), anti-PD-1-BV421 (BioLegend), anti-CXCR5-AF488 (BD Biosciences), anti-CD45RA-A700 (BioLegend), anti-CCR7-PerCPcy5.5 (BioLegend).

All cells were sorted for ATC-Seq and RNA-Seq using a BD FACSAria. Gating strategies for sorted subsets were as follows: fCD8; CD3^+^CD4^-^CD8^+^CD45RA^-^ CXCR5^+^, non-fCD8; CD3^+^CD4^-^CD8^+^CD45RA^-^CXCR5^-^, Naïve CD8^+^ T cells; CD3^+^CD4^-^CD8^+^CD45RA^+^CCR7^+^, GCTfh; CD3^+^CD4^+^CD8^-^PD-1^high^CXCR5^high^, non-Tfh; CD3^+^CD4^+^CD8^-^PD-1^-^CXCR5^-^. For RNA-Seq, cell subsets were sorted in RLT buffer (Invitrogen) containing 1% beta-mercaptoethanol. For ATAC-Seq, cell subsets were sorted in PBS containing 5% fetal calf serum (FCS) for downstream processing. In all sorting experiments, the grade purity on the sorted cells was >95%.

### Immunofluorescence staining

Localization of CD8^+^ T cell subsets was assessed as described by (6). Briefly, slides were prepared from 4 μm sections of paraffin-embedded tissue blocks and immunostained using in-house optimized protocols. For each LN, serial sections were stained singly with antibodies against BCL6 and CD8 and a DAB DAB visualization kit (Envision Double Stain system, Dako; USA) for bright field microscopy. Alternatively, we used the Opal 4-Color Fluorescent IHC Kit (PerkinElmer, USA) for immunofluorescence microscopy light. Slides were mounted and viewed using the Axio observer and TissueFAXS imaging software (TissueGnostics). Quantitative imaging analysis was conducted with TissueQuest (TissueGnostics). Medians of the cell density in the scanned GCs were used to perform statistical analysis.

### DNA methylation and drug treatment assays

Specific CpG within the *CXCR5* gene region was measured for DNA methylation according to a protocol from Paulin et al., (55). Briefly, a minimum of 500 ng of genomic DNA was bisulfide treated and amplified using a primer designed to cover 500 bp around the TSS. Amplified product was then analysed using Agena MassArray platform.

Drug treatment was then performed on the same samples used for DNA methylation assay. Briefly, an average of 100,000 non-fCD8s were sorted from the lymph node tissues and treated for 24 hrs with 10 μM of 5’-aza-2-deoxycytidine; a drug that inhibits the activity of genome-wide methyl-transferases. Thereafter, cells were washed, lysed and RNA were extracted and purified. cDNA was generated from the purified RNA using (Bio-Rad). CXCR5 mRNA transcripts were then measured from the generated cDNA using digital droplet PCR (ddPCR).

### Chemotaxis assay

Chemotaxis assays were performed as previously described (56). Briefly, LNMCs were suspended at a density of 1 × 10^6^ in RPMI 1640 medium containing L-glutamine, antibiotics, 10 mM HEPES buffer and 0.5% fatty acid-free BSA. Cells were cultured for 30-60 min at 37°C before being plated in trans-well inserts with a pore size of 5 μm and a diameter of 6.5 mm in 24-well plates (Corning Costar). 100 ul cells (1 × 10^6^) were added to the upper wells and 580 ul diluted CXCL13 chemokine (Peprotech) at 50 ng/ml was placed in the bottom wells, and plates were incubated for 3 hours at 37°C in 5% CO_2_. Migrated cells were stained with viability dye, CD3, CD4, CD8, CXCR5, and PD-1, and counted using flow cytometry.

### ATAC-Seq

Library preparations were performed as described by (39). Briefly, an average of 20,000 cells was sorted from LNs for fCD8s, non-fCD8s, naïve CD8^+^ T cells, GCTfh and non-Tfh. Five biological replicates were sorted for each subset. Sorted cells were lysed using lysis buffer (10mM Tris-HCL, pH 7.4, 10mM NaCl, 3mM MgCl_2_, 0.1% IGEPAL CA-630). Lysed cells were treated with 2.5 μl of Tn5 Transposase (Illumina, San Diego, CA) suspended in 50 μl of 1X TD buffer for 30 minutes at 37°C. Thereafter, transposed DNA was purified using QiaQuick MiniElute columns (Qiagen, Valencia, CA). Purified transposed DNA was amplified by PCR using Nextera barcoded primers (Illumina, San Diego, CA) and NEBNext High-Fidelity 2X PCR Master mix (New England Biolabs) with 12 cycles. Barcoded amplified libraries were purified using QiaQuick MiniElute columns (Qiagen, Valencia, CA) and quantified with KAPA real-time library quantification kit (KAPA, Wilmington, Massachusetts). Paired-end sequencing was performed on the high throughput NextSeq 500 (Illumina, San Diego, CA). Raw data from sequencer were stored in an on-onsite database and is available on request.

### RNA-Seq

An average of 20,000 cells were sorted directly into lysis (RLT) buffer (Qiagen, Valencia, CA) for RNA-Seq. Subsets that were sorted are: fCD8s, non-fCD8s, naïve CD8^+^ T cells, GCTfh and non-Tfh. Five biological replicates were used to perform this experiment. Total RNA was isolated from lysed cells using Qiagen RNeasy Mini columns (Qiagen, Valencia, CA) according to the manufacturer’s instructions. Purified RNA was evaluated with BioAnalyzer RNA pico kit (Agilent Technologies Inc, Santa Clara, CA). Messenger RNA (mRNA) was isolated from total RNA using NEBNext oligo dT beads (New England Biolabs). Isolated mRNA was fragmented and thereafter reverse transcribed to cDNA using NEBNext ultra RNA library preparation kit (New England Biolabs). The cDNA products were purified using AmpureXP beads (Beckman Coulter, Danvers, MA) and indexed using NEBNext multiplex oligo (New England Biolabs). Size distribution was evaluated using Agilent high-sensitivity DNA chip and initial quantification was performed using Qubit dsDNA high sensitive kit (ThermoFisher Scientist, Waltham, MA) and the median obtained on the Tapestation (Agilent Technologies Inc). KAPA kit was used for final quantification of obtained cDNA libraries molarity for sequencing. Index libraries were pooled and sequenced using high throughput NextSeq 500 (Illumina, San Diego, CA). Raw data from sequencer was stored in an on-onsite database and is available on request.

### Statistical analysis

Statistical analyses were conducted using Prism software, version 6.0 (GraphPad, Inc.). Two-tailed tests were employed, and p-values less than 0.05 were considered significant. Analysis on the next generation sequencing data is described in the Bioinformatics analysis below.

### ATAC-Seq analysis

To detect open chromatin regions (OCR) ATAC-seq Illumina reads were first filtered and trimmed for quality using TrimGalore and passed through the Kundaje lab pipeline (57) that performed the necessary quality controls (filtering of duplicate reads, removing reads mapping to the mitochondria) and peak detection together with irreproducible discovery rate (IDR) analysis using the biological replicates for each cell type. A cutoff of 0.1 was chosen for IDR. An optimal set of peaks that was produced for each cell type by the Kundaje pipeline was used for downstream analysis. OCR regions were compared between cell types using the DiffBind and EdgeR (58). A cut-off of 0.05 was chosen for FDR. We calculated the differential OCR using only the cell subsets as contrasts and subsequently paired the samples according to the patient from which the cells were extracted. The second method proved to be more sensitive at the same FDR of 0.05. PCA was performed using the top 1000 OCR by variance. The same sites were also used to construct a heatmap using the dba.heatMap function. Peak regions were annotated with the annotatePeak function from the ChIPseeker package (59). Annotations further than 50kb upstream from the TSS or those 10kb beyond the 3’-end of the gene were excluded. Gene ontology (GO) term enrichment was calculated with the enrichGO function from clusterProfiler (59).

### RNA-Seq analysis

RNA-Seq short reads were quantified using Kallisto (60). The Ensembl version 85 (GRCh37) was used as a transcriptome reference. Options were included to correct for “GC bias” and bootstrap sampling of 100. The Sleuth R package was used for downstream quantification and differential expression analysis (61). Gene transcripts were aggregated to gene level using internal sleuth functions. When doing pairwise comparisons (e.g. fCD8 vs non-fCD8), the design matrix was constructed in a way that would take the natural variation of expression data between subjects into account. Thus, the reduced design formula took the shape of *~pid*, while the full model *~pid + condition*, where pid refers to the patient id and condition refers to the cell type. The likelihood ratio test (LRT) of Sleuth was used to determine differential expression of genes by determining whether the *condition* variable added significant contribution in explaining the count data. Additionally, to determine the effect size of differential gene expression, the beta value for the condition variable was used as a proxy for log-fold differences in gene expression between conditions. For visualization purposes, the batch effects introduced by individual patients were removed using the remove Batch Effects function of the R package limma. Functional enrichment was determined using both the enrichGO and gseGO functions of the clusterProfiler package.

### Transcription factor footprinting and enrichment

Wellington-bootstrap was used for footprint detection (43). To increase sensitivity of footprint prediction, aligned reads in the form of BAM files were merged for each cell type: fCD8, non-fCD8, Naive CD8^+^T cells, GCTfh. For the HIV-Specific cell sets, reads were not merged to determine HIV-Specific footprinting sets. Predicted footprints were extended by 5 bp at each end and TF matching was performed using RGT (62). We used both the HOCOMOCO (63) and JASPAR (64) databases to complement mutually exclusive transcription factors from each set, e.g. *Id2* is not included in JASPAR, but is included in HOCOMOCO. Predicted footprints were filtered if they were more than 50 kilobases upstream from the transcription start site. Transcription factors that did not have evidence of expression from the RNA-Seq data were also filtered. We determined TF enrichment by comparing the frequency of predicted TF motifs in footprints compared to a background random set generated by RGT using a Fisher exact test. FDR values were determined using the R package *qvalue* (65) and a cut-off of 0.01 was used to filter out non-significant hits. We contrasted subjects for differential enrichment of TF motifs detected within the predicted footprints. We used the Wellington Bootstrap method (42) to detect differential footprints that can indicate higher activity of a transcription factor at different footprint loci. Differential footprints were chosen on the criteria of having a score >8 as produced by the *wellington_bootstrap.py* script or if a footprint was exclusively detected in a condition.

We calculated the differentially enriched TF motifs between all the cell types, i.e. fCD8, non-fCD8, Naive CD8^+^ T cells, GCTfh, and each of the HIV-Specific sets, yielding 27 comparisons. For the fCD8 and non-fCD8 subsets, we compared the enrichment of TF footprints between up and down regulated genes. This was done for both the predicted footprints from the whole set as well as the footprints demonstrating differential signal produced by Wellington bootstrap. For the wellington bootstrap, relative frequencies of TF motifs were calculated. We then clustered these relative frequencies and displayed them as a heatmap.

Plots for the footprints were generated based on the average Tn5 insertion sites 200bp around the predicted footprinting sites. Because Tn5 does have cleavage bias, the counts were corrected using the *tracks* module of the RGT-HINT package. Additional plots were generated for differential footprints.

### Weighted correlation network analysis

We tested the modularity of gene expression using weighted correlation network analysis (WGCNA) (47). For the RNA-Seq data, raw count data was first regularized with the variance stabilizing transformation (vst) function from DESeq2 (66). WGCNA is sensitive to the amount of available data for network construction. We therefore included Naive CD8^+^ T cells, non-fCD8, fCD8, and GCTfh and a set of HIV specific samples (blood and LN) to augment our network for both the expression network (RNA-Seq data) as well as the chromatin accessibility network (ATAC-Seq data). After construction of the gene expression network, GSEA was performed to determine the level of enrichment of a module in a subset. For this, the data were adjusted to account for batch effects. For each gene, a Z-score was calculated and sorted and used as input for the GSEA and the results visualized.

To determine whether there are modules of OCR specific to expressed genes, a WGCNA network for the OCR regions from the ATAC-Seq data was constructed. We used the read counts from the merged peaks calculated by DiffBind as input and also regularized the input with variance stabilizing transformation. We hypothesized that while OCR in and around genes would be largely correlated, certain OCRs may be in different modules depending on the subset. To test this, we assigned all the OCRs to modules and then used the OCR annotation as a gene reference. We specifically looked at genes that are differentially expressed in fCD8 and GCTfh compared to non-fCD8 and cross-referenced this with the ATAC-Seq WGCNA network modules. Similarly to the WGCNA for RNA-Seq, we performed GSEA using batch adjusted count data and calculated a Z-score for each subset, ranked these values and used them as input for GSEA.

### Nucleosomal Positioning

NucleoATAC (67) was used to predict nucleosome occupancy and position from the ATAC-Seq data. For each subset, MACS 2 was used with the *--broadPeak* option to localize regions for nucleosomal detection. These regions were further expanded by 200bp on either end. To improve signal, samples reads were merged within each subset.

To investigate differences in nucleosomal positioning within the promoter region of CXCR5 between GCTfh and fCD8, the region matching the promoter of TSS was successively trimmed from the 3’ end. With each successive trim, NucleoATAC was again run on that region to calculate nucleosomal occupancy signals and positions. This trimming should bias the removed shorter reads and reveal temporal positioning of the nucleosome. Importantly, the fragment size distribution files and V-matrix files produced from the full peakset was used as input to eliminate fragment distribution bias, produce a BED file containing these overlapping regions. The smoothed signal was plotted and the combined position file was used for dyad positioning of the nucleosome.

## Acknowledgements

We thank our study participants, the laboratory staff at the HIV Pathogenesis Programme and the clinic staff at the Prince Mshiyeni Hospital, Durban, South Africa. We also thank Professor John E. Wherry, John L. Johnson and Sasikanth Manne at the Institute of Immunology, Philadelphia, PA, USA for training on ATAC-Seq and technical guidance on data analysis. Additionally, we thank Dr. Erica Andersen-Nissen and Professor Julie McElrath at the HIV Vaccine Trial Network (HVTN), Cape Town for access to the HVTN sequencing platform. We thank Dr. Daniel Muema for technical support on RNA-Seq library generation.

## Author contributions

ZMN conceived the project. ZNM and FJO designed the experiments. IJ and JP and ThanNg performed the lymph node biopsies. TNK processed the lymph node samples. FJO performed majority of the experiments described in this study under the supervision of ZMN with technical assistance from VR on drug treatment assay. OOB perform tissue staining and analysis with technical support from TRK. DNA methylation assay using bi-sulfite treatment and sequencing was performed by Inquaba Biotec. FJO and AFN performed the next generation sequencing (NGS). WS performed NGS analysis (ATAC-Seq and RNA-Seq) under the supervision of TO. FJO analysed other data reported in this study. FJO, WS and ZMN wrote the manuscript. ZMN, BDW and TN provided critical edits to the manuscript. All authors reviewed the manuscript.

## Corresponding authors

Address correspondence to Zaza M Ndhlovu

## Supplementary data

**Supplementary Figure 1A and B:** Study samples and experimental design for flowcytometry, tissue imgagingATAC-Seq, RNA-Seq and DNA methylation. Experimental setup describing cell subsets and markers used in cell sorting.

**Supplementary Figure 2:** Heatmap of up-regulated genes with epigenetic function in fCD8. The heatmap shows the relative rank of gene expression (after batch-adjustment) of the epigenetic genes. Majority of the genes are involved in histone modification as shown in the heatmap.

**Supplementary Figure 3: (A)** The heatmap shows a condensed overview of ATAC-Seq signal of the top 10% ATAC-Seq peaks by variance. The clusters are organized in a hierarchical fashion showing subset specific clusters. **(B)** The figure shows deferentially expressed genes with corresponding differential accessibility OCRs proximal to the gene. The y-axis represents the log_2_ fold change in gene expression, while the x-axis represents the log fold change in chromatin accessibility. A regularized regression line is fitted to the data. Example genes are annotated. The gene of interest, *CXCR5* is coloured in red.

**Supplementary Figure 4: (A)** We determined the top TF enriched using WB for each subset and plotted the results on pie chart. ATAC-Seq peaks 11kb from the TSS region of CXCR5 ATAC seq peaks within 11kb of the TSS of CXCR5 are shown. The boxes indicate the named upstream regions, i.e. U1 (−6.5kb) and U2 (−11kb). **(B)** Set enrichment of TF. We ranked ATAC-Seq signals of OCRs within the module 5 with the representative *eigengene* of the module 5. Regions were sorted in descending order depending on their correlation with the module 5. For each subset, we used the calculated TF footprints in each region and determined by *set enrichment analysis* whether these TFs were likely enriched in regions higher correlated with the module 5 eigengene. That is, we hypothesize TF showing higher SEA enrichment with the module 5 eigengene to be more associated with the hub regions that are purported to be central in governing accessibility programs across this module. TF were aggregated at family level. From the figures, it becomes apparent that there is a progressive enrichment of MAF-family related factors from non-fCD8s to the enrichment of pioneering POU-family transcription factors in fCD8s with *GCTfh* sharing these *TFs*. High enrichment is shown as positive (red) values, while negative enrichment (i.e. TF depleted module 5 OCRs) are shown in blue.

**Supplementary Figure 5:** Based on experimental and computational evidence generated in this study, we propose that in naïve CD8^+^ T cells, DNA methylation of CpG islands around the TSS stably silence *CXCR5* gene expression by attracting chromatin remodelling proteins and histone modifiers to the loci which compact chromatin around the TSS into heterochromatin state. Cell division following TCR stimulation results in partial chromatin relation and passive DNA demethylation around the promoter region allowing for basal transcriptional activity observed in non-fCD8s relative to naïve CD8^+^ T cells. As the cells continue to divide, a small proportion of cells become more extensively demethylated at the *CXCR5* gene loci and gradually accumulate epigenetic regulatory proteins including pioneer factors (the POUs), namely POU3F3 and POU3F1 which are recruited to the TSS proximal regions. These factors decondense the chromatin at the TSS thereby exposing unmethylated DNA for transcription, thus allowing the transcription machinery to bind and transcribe the *CXCR5* gene.

**Supplementary Figure 6: (A)** The animation (left) is a cartoon showing hypothesized translocation events in the TSS region of CXCR5. This figure was generated from data produced by NucleoATAC. At each iteration, short reads were progressively removed from the 3’→ 5’ end and a new nucleosomal signal generated by NucleoATAC. We observe a shift in nucleosomal positioning in both fCD8s (red) and GCTfh (blue), but a more pronounced depletion of nucleosomal signal close to the TSS of CXCR5 and subsequently a higher peak further upstream, whereas nucleosomal occupancy is determined to be mostly proximal to the TSS in fCD8s. On the right, a ARToon model is drawn depicting average counts of CXCR5 transcripts produced by each cell subset, with GCTfh quickly outpacing fCD8s. **(B)** Nucleosomal positioning can dictate transcription efficiency. We postulate that fCD8s have less CXCR5 expression relative to GCTfh due to higher nucleosomal occupancy around the CXCR5 TSS. The rationale is as follows, although, we detected primary nucleosomal signal over the TSS in both fCD8s and GCTfh, the secondary nucleosomal signal is closer to the TSS in fCD8s but further upstream in GCTfh. This suggest that the repositioning of the nucleosome further away from the TSS, in GCTfh, makes it easier for the transcriptional machinery to access the promoter and initiate transcription. Nucleosomes are pushed away from gene promoter regions by a family of proteins called nucloesomal remodellers. Some remodellers are more efficient at evicting nucleosomes from active gene loci than others (68). Interestingly, fCD8s and GCTfh express different types of nucloesomal remodellers. Therefore, we postulate that nucleosomal remodellers in GCTfh are more efficient at pushing the nucleosome further upstream, which completely uncovers the CXCR5 TSS for transcription whereas, fCD8s nucleosomal remodellers are less efficient at pushing the nucleosome away from the TSS, hence the attenuated *CXCR5* gene expression.

**Supplementary data file:** List of differentially expressed genes between fCD8s and non-fCD8s. Top 285 genes highlighted in red are upregulated in fCD8s while the bottom 322 genes highlighted in green are downregulated in fCD8s compared to non-fCD8s.

